# Characterization of human immunodeficiency virus (HIV-1) envelope glycoprotein variants selected for resistance to a CD4-mimetic compound

**DOI:** 10.1101/2022.04.21.489129

**Authors:** Saumya Anang, Jonathan Richard, Catherine Bourassa, Guillaume Goyette, Ta-Jung Chiu, Hung-Ching Chen, Amos B. Smith, Navid Madani, Andrés Finzi, Joseph Sodroski

## Abstract

Binding to host cell receptors, CD4 and CCR5/CXCR4, triggers conformational changes in the human immunodeficiency virus (HIV-1) envelope glycoprotein (Env) trimer that promote virus entry. CD4 binding allows the gp120 exterior Env to bind CCR5/CXCR4 and induces a pre-hairpin intermediate conformation in the gp41 transmembrane Env. Small-molecule CD4-mimetic compounds (CD4mcs) bind within the conserved Phe-43 cavity of gp120, near the binding site for CD4. CD4mcs inhibit HIV-1 infection by competing with CD4 and by prematurely activating Env, leading to irreversible inactivation. BNM-III-170 is a CD4mc that inhibits the infection of approximately 70% of HIV-1 strains at micromolar concentrations. We selected and analyzed variants of the primary HIV-1_AD8_ strain resistant to BNM-III-170. Two changes (S375N and I424T) in gp120 residues that flank the Phe-43 cavity each conferred ∼5-fold resistance to BNM-III-170 with minimal fitness cost. A third change (E64G) in Layer 1 of the gp120 inner domain resulted in ∼100-fold resistance to BNM-III-170, ∼2-3-fold resistance to soluble CD4-Ig, and a moderate decrease in viral fitness. The gp120 changes additively or synergistically contributed to BNM-III-170 resistance. The sensitivity of the Env variants to BNM-III-170 inhibition of virus entry correlated with their sensitivity to BNM-III-170- induced Env activation and shedding of gp120. The S375N and I424T changes, but not the E64G change, conferred resistance to BMS-806, a potent HIV-1 entry inhibitor that blocks Env conformational transitions. These studies identify pathways whereby HIV-1 can develop resistance to CD4mcs and BMS-806 conformational blockers, two classes of entry inhibitors that target the conserved gp120 Phe-43 cavity.

**IMPORTANCE:** CD4-mimetic compounds (CD4mcs) and BMS-806 are small-molecule inhibitors of human immunodeficiency virus (HIV-1) entry into host cells. Although CD4mcs and BMS-806 inhibit HIV-1 entry by different mechanisms, they both target a pocket on the viral envelope glycoprotein (Env) spike that is used for binding to the receptor, CD4, and is highly conserved among HIV-1 strains. Our study identifies changes near this pocket that can confer various levels of resistance to the antiviral effects of both a CD4mc and BMS-806. We relate the antiviral potency of a CD4mc against this panel of HIV-1 variants to the ability of the CD4mc to activate changes in Env conformation and to induce the shedding of the gp120 exterior Env from the spike. These findings will guide efforts to improve the potency and breadth of small-molecule HIV-1 entry inhibitors.

## INTRODUCTION

The binding of the human immunodeficiency virus (HIV-1) envelope glycoprotein (Env) to host cell receptors, CD4 and CCR5/CXCR4, triggers virus entry into the cell (1–11). The Env trimer consists of three gp120 exterior Envs and three gp41 transmembrane Envs. Prior to receptor engagement, the HIV-1 Env trimer on virions can potentially sample three conformations, a pretriggered “closed” conformation (State 1), the “open” CD4-bound conformation (State 3) and an intermediate “partially open” conformation (State 2) (1, 12–15). Primary HIV-1 Envs are maintained in State 1 to various degrees by multiple intramolecular and intersubunit interactions, rendering these Envs relatively resistant to the binding of potentially neutralizing antibodies. CD4 binding drives Env from State 1 to State 2 and then into State 3, the prehairpin intermediate (12–18). In the prehairpin intermediate, the heptad repeat (HR1) region of gp41 forms an exposed coiled coil (16–19). Binding of the State 3 Env to the CCR5 or CXCR4 coreceptor is thought to induce the formation of a highly stable gp41 six-helix bundle, which promotes the fusion of the viral and cell membranes (20–25).

The surface of the HIV-1 Env trimer exhibits significant strain variability and heavy glycosylation, which contributes to the avoidance of potentially neutralizing antibodies (26–29). The binding site for CD4 on gp120 consists of a conserved surface that is conformationally altered by CD4 binding. CD4 binding creates an internal pocket in gp120 called the Phe-43 cavity that is bounded by highly conserved residues from gp120 and a single phenylalanine residue (Phe 43) from CD4 (30). The ∼150-Å^3^ Phe-43 cavity is a target for two classes of small-molecule HIV-1 entry inhibitors, the CD4- mimetic compounds (CD4mcs) and conformational blockers like BMS-806 (30–40).

CD4-mimetic compounds (CD4mcs) disrupt HIV-1 entry by binding to gp120 in the Phe-43 cavity, directly competing with CD4 but also prematurely triggering Env (31, 40–42). Thus, CD4mcs drive the State-1 Env trimer into downstream conformations (States 2 and 3) that, in proximity to a coreceptor-expressing target cell, can mediate HIV-1 infection (42). These CD4mc-induced Env intermediates are short-lived and, in the absence of a coreceptor-expressing target cell, apparently decay into inactive, dead-end conformations (41, 42). At CD4mc concentrations that do not completely inhibit HIV-1 infection, the induction of State-2 or State-3 Env conformations sensitizes HIV-1 viruses to neutralization by otherwise poorly neutralizing antibodies (43, 44).

Such antibodies recognize epitopes induced by CD4, including the V2 and V3 variable regions of gp120, as well as the highly conserved gp120 region that overlaps the coreceptor-binding site (45).

The CD4-bound conformation of the HIV-1 Env on the cell surface can also be targeted by antibody-dependent cell-mediated cytotoxicity (ADCC) (46–48). In HIV-1- infected cells, complexes of Env and CD4 can be recognized by antibodies like 17b against the conserved CD4-induced (CD4i) gp120 region overlapping the coreceptor-binding site (49–52). Binding of CD4i antibodies promotes binding of antibodies like A32 against the highly conserved “Cluster A” epitopes in the gp120 inner domain (53).

These antibodies cooperate to induce the asymmetric State 2A Env conformation, which is efficiently targeted by ADCC (54). The HIV-1 Nef and Vpu proteins decrease the expression of Env-CD4 complexes on the infected cell surface, reducing the vulnerability of infected cells to ADCC (55–61). CD4mcs induce CD4i conformations of Env but, unlike CD4, are not susceptible to downregulation by Vpu or Nef (62). Thus, CD4mc can sensitize HIV-1-infected cells to ADCC by CD4i and anti-Cluster A antibodies that are present at high titer in most HIV-1-infected individuals (49, 62–67). CD4mcs are being evaluated in animal models for their ability to stimulate ADCC as a means to reduce the size of the infected cell reservoir (68).

CD4mcs are also being evaluated for their ability to prevent acquisition of HIV-1.

Intravaginal application of a CD4mc, either before or simultaneously with virus challenge, protected bone marrow-liver-thymus humanized mice from HIV-1 infection (69). Although the development of an effective HIV-1 vaccine has been frustrated by the inefficient elicitation of broadly neutralizing antibodies, antibodies that neutralize HIV-1 sensitized by a CD4mc are readily elicited by Env immunogens (43, 44, 62, 63) . Antibodies elicited by monomeric gp120 protect monkeys from stringent, heterologous simian-human immunodeficiency virus (SHIV) mucosal challenge, provided the challenge virus is exposed to a CD4mc (43).

Early CD4mcs discovered using a gp120-CD4 screen exhibited weak antiviral potency against a limited range of HIV-1 isolates (31, 40). Iterative cycles of design, guided by CD4mc-gp120 structures and empirical testing, have led to the development of analogues with improved potency (36-38, 70-72). Primary HIV-1 strains exhibit a range of sensitivity to the improved CD4mcs. Progressive increases in CD4mc potency have been accompanied by an increase in the breadth of activity against a wider range of HIV-1 strains (37, 38). BNM-III-170 is currently the most potent and well-studied CD4mc, and inhibits approximately 70% of a global panel of multi-clade HIV-1 variants (37).

Despite significant improvements in the potency and breadth of CD4mcs, some primary HIV-1 strains remain resistant to their antiviral effects (37). The basis for resistance to the CD4mcs is two-fold. First, although the gp120 residues lining the Phe- 43 cavity are well conserved among HIV-1 strains, some variation is tolerated. HIV-1 strains that are Clade AE recombinants have a histidine residue (His 375) that fills the Phe-43 cavity, preventing efficient binding of CD4mcs (73–75). For most other HIV-1 strains, the Phe-43 cavity is not filled and is therefore hypothetically available for binding of the CD4mc (74, 75). Except for Ser 375, the gp120 residues lining the Phe 43-cavity are conserved among the non-Clade AE HIV-1 (74, 75). Ser 375 is occasionally replaced by a threonine residue, and much less commonly by isoleucine and methionine residues. Viruses with Thr 375 are even more sensitive to CD4mcs than the corresponding viruses with Ser 375 (73, 76). By contrast, S375I and S375M HIV-1 mutants are more resistant to inhibition by the CD4mcs. Thus, only in a small subset of naturally occurring HIV-1 strains does variation in the gp120 residues lining the Phe-43 cavity account for resistance to the CD4mcs.

A second mechanism of resistance is based on the requirement that to bind and inhibit HIV-1, CD4mcs must induce transitions from State 1 to downstream conformations (26, 31, 35–37, 41, 42, 53, 54, 70–72, 77). Primary HIV-1 strains exhibit a continuous range of “triggerability,” a property that is inversely related to the height of the activation barrier separating State 1 and State 2 (13, 26, 27). Viruses with Envs that have stable State-1 conformations and therefore are less prone to make transitions from State 1 are expected to exhibit greater resistance to CD4mcs (26, 78–81). This second mechanism of resistance is more common than alterations within the Phe-43 cavity and reflects the different levels of responsiveness to CD4 that are advantageous to HIV-1 in different natural circumstances. Thus, the rank order of susceptibility of HIV-1 strains to CD4mcs remains relatively constant, even as the potency of CD4mc analogues increases (37).

To investigate pathways to CD4mc resistance preferred by HIV-1 in the absence of immune selection, we passaged the primary HIV-1_AD8_ strain in the presence of increasing concentrations of BNM-III-170. We identified three changes in gp120 that contribute individually and additively to the resistance of the selected viruses. We evaluated the relative impact of these gp120 changes on CD4 binding and viral fitness. Some of the changes associated with BNM-III-170 resistance also contributed to resistance to BMS-806, another small-molecule entry inhibitor that shares part of its gp120 binding site with that of the CD4mcs (33-37, 82-86). We examined the sensitivity of the BNM-III-170-resistant viruses to conformation-dependent Env ligands; the results indicate that BNM-III-170 escape in this case is determined by local changes rather than global effects on Env conformation. These results provide valuable information about HIV-1 pathways to achieve CD4mc resistance and can guide future efforts to develop more potent and broad CD4mcs.

## RESULTS

### Selection of BNM-III-170-resistant HIV-1_AD8_

We sought to evaluate natural pathways that HIV-1 can utilize to achieve resistance to a CD4mc, BNM-III-170, while maintaining replication competence. To this end, we passaged molecularly cloned HIV-1_AD8_ in C8166-R5 cells in the presence of increasing concentrations of BNM-III-170; starting with 5 µM BNM-III-170, the concentration of the CD4mc was gradually increased to 130 µM until no further effects on virus replication were observed (Fig. 1A). For the first 28 days, the BNM-III-170-treated infected cells produced less virus in the culture medium than the untreated infected cells. The reverse transcriptase levels in the BNM-III-170-treated cultures were higher than those in mock-infected cells, suggesting that some level of HIV-1_AD8_ replication still occurred in the presence of the CD4mc. Virus levels in the medium of the BNM-III-170-treated infected cells increased after day 28, approaching but not reaching the levels in the untreated infected cultures.

**Figure 1.**
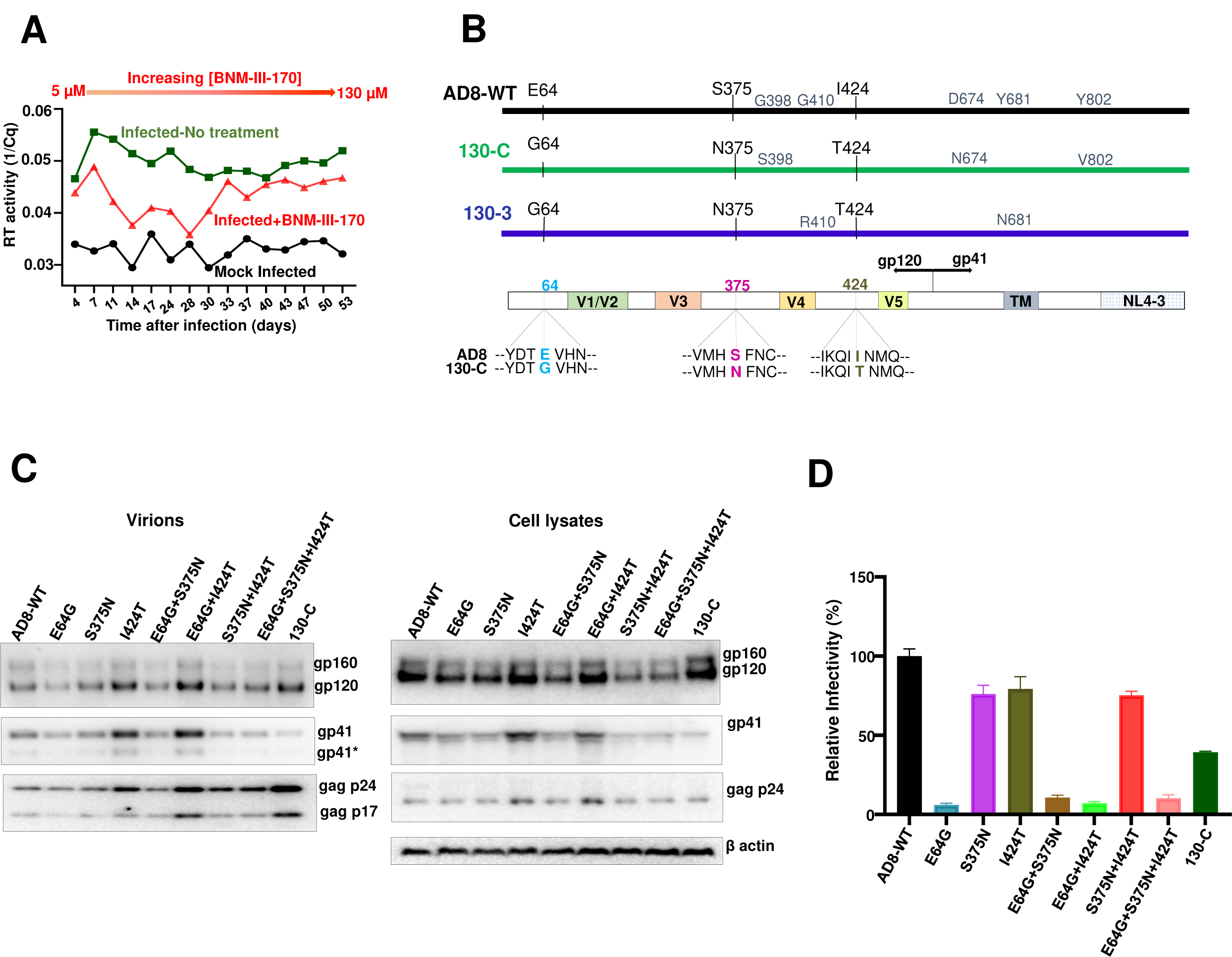
Selection and characterization of BNM-III-170-resistant HIV-1_AD8._ (A) The replication of HIV-1_AD8_ in C8166-R5 cells in the absence and presence of increasing concentrations (5-130 µM) of BNM-III-170 is shown. The results from mock-infected cultures are also shown. Virus produced from HEK 293T cells transfected with the pNL4-3-AD8 proviral construct was incubated with C8166-R5 cells. Virus replication was monitored by measuring reverse transcriptase in the culture medium by quantitative real-time PCR. (B) A schematic representation of the wild-type (WT) HIV-1_AD8_ Env is shown, with the changes in two different cloned Envs (130-C and 130-3) derived from the viruses from the BNM-III-170-treated cells. In the bottom diagram, the changes common to both cloned Envs are shown. The V1/V2, V3, V4 and V5 variable gp120 regions, gp120/gp41 cleavage site and transmembrane region (TM) are shown. The gp41 cytoplasmic tail of the virus is a chimera of HIV-1_AD8_ and HIV-1_NL4-3_ Env sequences. (C) The expression level, processing and virion incorporation of the indicated Env variants are shown. HEK 293T cells were transfected with pNL4-3-AD8 proviral constructs expressing the Env variants. Cell lysates and viruses were harvested 48 h later and analyzed by Western blotting for the indicated proteins. A minor form of gp41 resulting from cleavage of the cytoplasmic tail (114) is designated by an asterisk. (D) The infectivity of viruses with the indicated Envs was measured on Cf2Th-CD4/CCR5 cells. HEK 293T cells were transfected with plasmids encoding the indicated Envs and HIV-1 packaging proteins and a luciferase-expressing HIV-1 vector. Forty-eight h later, cell supernatants were cleared and added to Cf2Th-CD4/CCR5 cells. Two days later, the cells were lysed and the luciferase activity was measured. The relative infectivity of viruses with the mutant Envs is compared with that of viruses with wild-type (WT) HIV-1_AD8_ Env. The results shown are representative of those obtained in three or more independent experiments.

To characterize HIV-1_AD8_ Env variants that were potentially selected during passage in the presence of BNM-III-170, *env* DNA was amplified by polymerase chain reaction and sequenced. The predicted amino acid sequences of two Envs (130-C and 130-3) from the BNM-III-170-treated culture are compared with that of the parental HIV-1_AD8_ Env in Fig. 1B. Three amino acid changes (E64G, S375N and I424T) were found in both Envs derived from the BNM-III-170-treated culture. All three changes in the gp120 glycoprotein are unusual in natural HIV-1 strains (74, 75). Glutamic acid 64 is nearly invariant in HIV-1 strains. Approximately 74% of HIV-1 strains have a serine residue at 375, and the remainder have threonine, histidine, isoleucine or methionine residues at this position; Asn 375 is very rare in natural HIV-1 strains. Isoleucine (46%) and valine (54%) predominate at residue 424 in natural HIV-1 variants; Thr 424 is highly unusual.

The three gp120 changes associated with BNM-III-170 selection were introduced individually or in combination into the HIV-1_AD8_ Env. The processing and incorporation into virion particles of these Env variants were comparable to those of the wild-type (WT) HIV-1_AD8_ and 130-C Envs (Fig. 1C). All Env variants detectably supported the entry of pseudotyped viruses into Cf2Th-CD4/CCR5 cells, although the infectivity of the viruses with the E64G change was significantly lower than that of WT HIV-1_AD8_ (Fig. 1D). Apparently, the E64G change in Env decreases the fitness of HIV-1, whereas the S375N and I424T changes are functionally well tolerated.

### Effects of Env changes on resistance to HIV-1 entry inhibitors

We examined the sensitivity of viruses pseudotyped with the Env variants to inhibition by BNM-III-170 and other gp120-directed entry blockers (Fig. 2 and Table 1). Viruses with the 130-C and 130-3 Envs were completely resistant to high concentrations (300 µM) of BNM-III-170, whereas the WT HIV-1_AD8_ was inhibited with a 50% inhibitory concentration (IC_50_) of 0.3 µM (Fig. 2A, left panel). Approximately 5-fold higher concentrations of BNM-III-170 were required to inhibit the S375N and I424T viruses (Fig. 2A, middle panel). Viruses with the E64G Env exhibited even greater (approximately 100-fold) resistance to BNM-III-170. In combination, the gp120 amino acid changes contributed additively to BNM-III-170 resistance (Fig. 2A, right panel).

**Figure 2.**
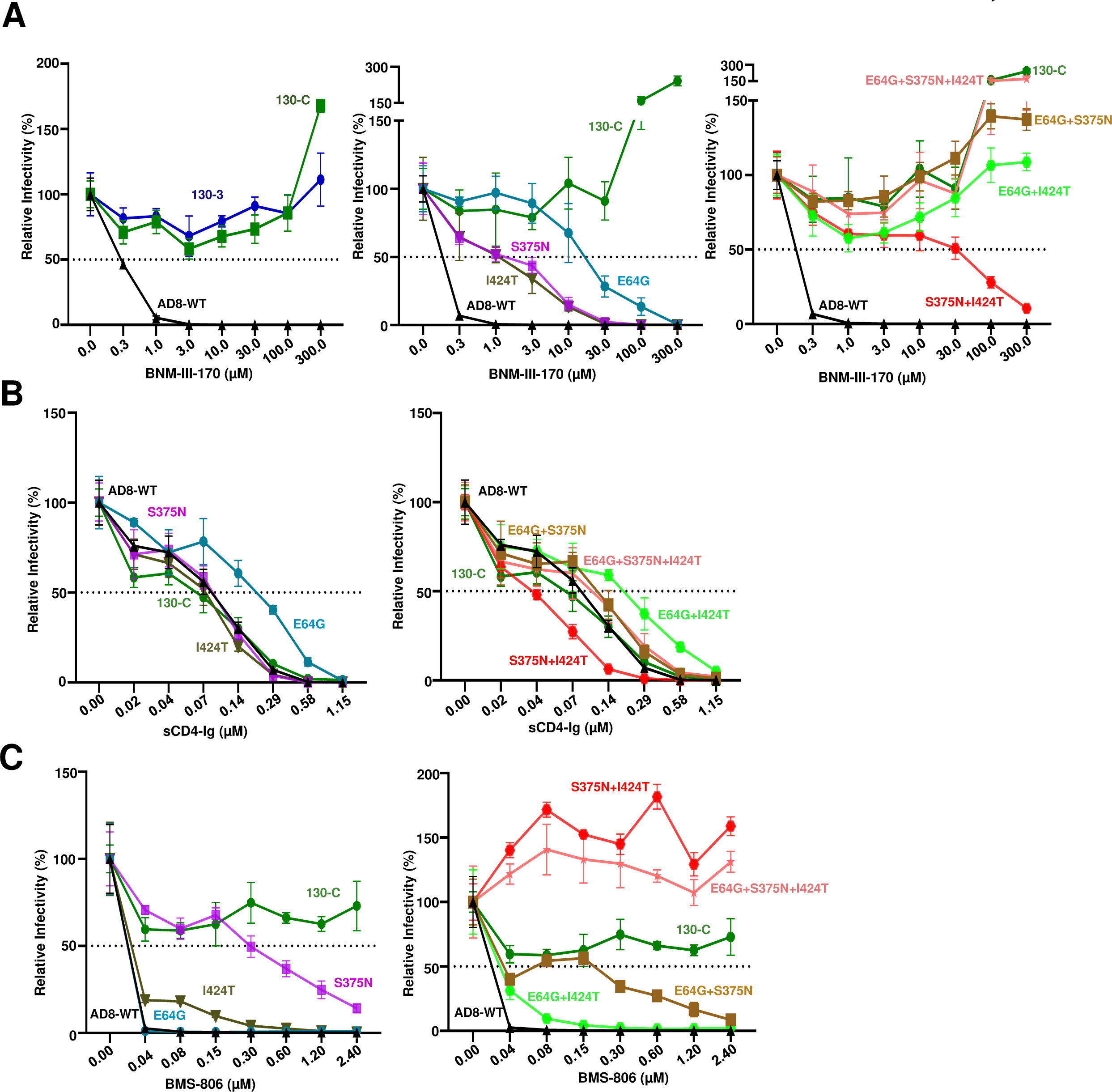
Effect of changes in HIV-1_AD8_ Env on virus sensitivity to BNM-III-170, sCD4-Ig and BMS-806. Recombinant luciferase-expressing viruses pseudotyped with the indicated Envs were produced as described in the Figure 1D legend. Viruses were incubated with the indicated concentrations of BNM-III-170 (A), sCD4-Ig (B) and BMS- 806 (C) for one h at 37°C. The viruses were then added to Cf2Th-CD4/CCR5 cells. After two days of culture, the cells were lysed and the luciferase activity was measured. The relative infectivity represents the luciferase activity obtained at each concentration of inhibitor compared with that obtained in the absence of inhibitor. The results shown are representative of those obtained in three or more independent experiments .

**Table 1.** Sensitivity of HIV-1_AD8_ Env variants to inhibition by BNM-III-170, sCD4-Ig and BMS-806. The sensitivity of viruses pseudotyped with the WT HIV-1_AD8_ and variant Envs to inhibition by BNM-III-170, sCD4-Ig and BMS-806 was determined as described in the Figure 2 legend. The IC_50_ values for each inhibitor are reported as means and standard deviations derived from at least two independent experiments.

| **IC<sub>50</sub>(μM)** | **BNM-III-170** | **sCD4-Ig** | **BMS-806** |
| --- | --- | --- | --- |
| **AD8-WT** | 0.23 ± 0.07 | 0.10 ± 0.08 | 0.02 ± 0.01 |
| **E64G** | 25 ± 3.1 | 0.23 ± 0.19 | 0.02 ± 0.01 |
| **S375N** | 1.2 ± 0.09 | 0.10 ± 0.09 | 0.30 ± 0.11 |
| **I424T** | 1.2 ± 0.08 | 0.08 ± 0.07 | 0.03 ± 0.01 |
| **E64G+S375N** | >300 | 0.12 ± 0.80 | 0.20 ± 0.1 |
| **E64G+I424T** | >300 | 0.21 ± 0.16 | 0.03 ± 0.01 |
| **S375N+I424T** | 48 ± 4.92 | 0.04 ± 0.02 | >300 |
| **E64G+S375N+I424T** | >300 | 0.10 ± 0.07 | >300 |
| **130-C** | >300 | 0.07±0.07 | >300 |

Viruses with the S375N + I424T changes could still be inhibited at high concentrations of BNM-III-170, whereas viruses with E64G in combination with either S375N or I424T were resistant to the CD4mc. Viruses with all three Env changes were as resistant to BNM-III-170 as viruses with the 130-C Env. We conclude that all three gp120 changes contribute to BNM-III-170 resistance, with E64G exerting the largest effect.

We examined the sensitivity of viruses pseudotyped by the HIV-1_AD8_ Env variants to soluble CD4-Ig (sCD4-Ig). Most of the variants, including 130-C, were inhibited by sCD4-Ig comparably to WT HIV-1_AD8_ (Fig. 2B and Table 1). The E64G and E64G + I424T viruses required 2-3-fold more sCD4-Ig for neutralization than the WT HIV-1_AD8_.

The S375N + I424T virus was neutralized by sCD4-Ig slightly more efficiently than WT HIV-1_AD8_. Thus, in the context of some Envs, slight decreases or increases in sensitivity to sCD4-Ig were associated with the E64G and S375N changes, respectively. Overall, small differences in the sensitivity of these variant viruses to sCD4-Ig were observed, relative to the profound differences seen for BNM-III-170.

BMS-806 is a potent small-molecule entry inhibitor that binds the pretriggered (State-1) conformation of Env and blocks CD4-induced conformational changes required for HIV-1 entry (12, 15, 19, 32, 83–86). The gp120 binding site of BMS-806 overlaps that of BNM-III-170 and other CD4mcs (33–37, 82) (see below). We evaluated the effect of the gp120 changes implicated in BNM-III-170 resistance on virus sensitivity to BMS-806. The WT HIV-1_AD8_ was inhibited completely at a BMS-806 concentration of 40 nM, whereas the 130-C virus was not inhibited even at a BMS-806 concentration of 2.4 µM (Fig. 2C and Table 1). The S375N change alone resulted in an approximately 15-fold reduction in sensitivity to BMS-806. A slight decrease in BMS-806 sensitivity was associated with the I424T change, whereas the E64G change had no effect on virus inhibition by BMS-806. The combination of S375N + I424T changes was sufficient to account for the full resistance of the 130-C virus to BMS-806.

### Env structures

We located the resistance-associated changes on structures of HIV-1 Env bound to the relevant ligands. In the process of binding CD4, gp120 components (the inner domain, outer domain and bridging sheet) are thought to rearrange around the internal Phe-43 cavity (30). Both Asn 375 and Thr 424 line the Phe-43 cavity, but do not directly contact CD4 in structures of gp120-CD4 complexes (Fig. 3A) (30). In the gp120-CD4 complex, Gly 64 is located in “Layer 1” of the gp120 inner domain and is distant from CD4 (87).

**Figure 3.**
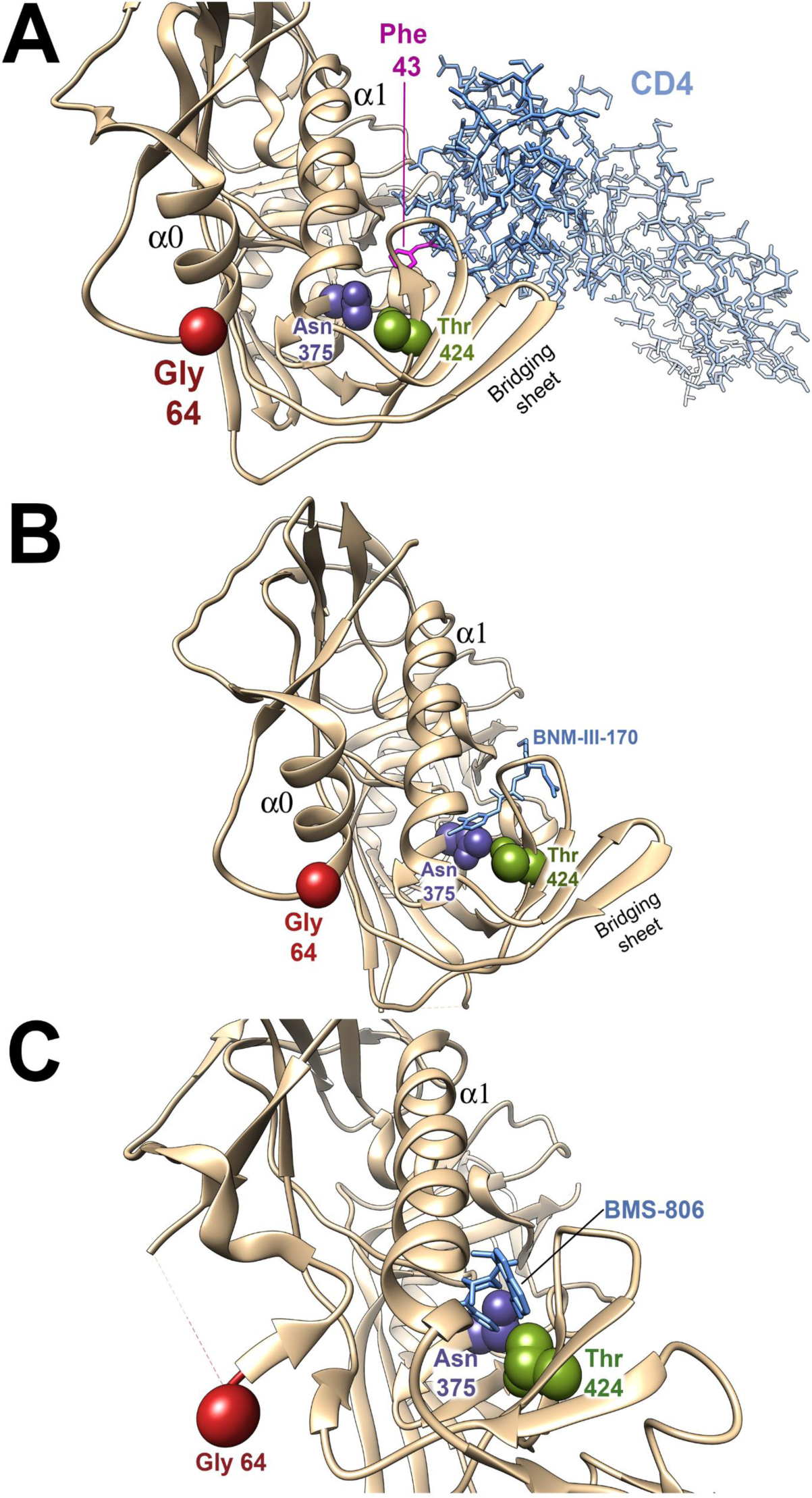
Location of amino acid residues implicated in virus resistance in Env structural models. The gp120 amino acid residues implicated in resistance to the inhibitors are shown on structures of HIV-1 Env-inhibitor complexes. In all structures, the resistance-associated changes have been introduced by the swapaa function in Chimera (115). (A) The HIV-1 gp120 core with N/C termini (tan ribbon) complexed with two-domain CD4 (light blue sticks) (PDB 3JWO) is shown (87). Phe 43 of CD4 is colored magenta. (B) The HIV-1 gp120 core with N/C termini (tan ribbon) complexed with BNM-III-170 (light blue sticks) (PDB 5F4P) is shown (87). (C) The HIV-1 sgp140 SOSIP.664 trimer (tan ribbon) complexed with BMS-806 (light blue sticks) (PDB 6MTJ) is shown (82). The Envs are oriented with the viral membrane at the top of the figure and the target cell membrane at the bottom. The α0 and α1 helices of gp120 are shown. Note that in C, the α0 helix is not present. Part of the gp120 Layer 1 loop between residues 59 and 64 is disordered in 6MTJ, and therefore the location of Gly 64 is an approximation.

During the process of gp120 binding to CD4, the formation of the α0 helix in Layer 1 allows His 66 and Trp 69 of gp120 to associate with the back of the Phe-43 cavity, decreasing the off-rate of CD4 (80, 87). The alteration of the highly conserved Glu 64 to glycine may decrease the efficiency with which the α0 helix forms.

In contrast to CD4, CD4-mimetic compounds like BNM-III-170 penetrate into the Phe-43 cavity (Fig. 3B) (35–37, 77). The S375N and I424T changes are expected to alter the shape of the Phe-43 cavity, explaining their individual and additive effects on BNM-III-170 antiviral potency. Although Gly 64 cannot contact BNM-III-170, the E64G change potentially decreases the Layer 1-Phe-43 cavity interaction, as discussed above for CD4. This interaction is also important for the binding of CD4mcs; for example, changes in His 66 in Layer 1 have been shown to decrease the antiviral efficacy of CD4mcs and sCD4 (78).

BMS-806 binds HIV-1 gp120 by occupying the Phe-43 cavity and adjacent water-filled channel (near Thr 424) (Fig. 3C) (34, 79). The alterations in the shape of these gp120 structures resulting from the S375N and I424T changes likely explain the observed decreases in BMS-806 potency. Gly 64 is distant from the BMS-806 binding site and since the formation of the α0 helix is not critical for BMS-806 binding (34, 82), the E64G change exerts little effect on its antiviral activity.

### Mechanisms of escape from BNM-III-170

The CD4-bound conformation of the HIV-1 Env trimer is more susceptible to disruption, resulting in the shedding of gp120 (88). We examined the effect of the Env changes on susceptibility of the virion spike to BNM-III-170-induced gp120 shedding at 37°C. The WT HIV-1_AD8_ Env shed nearly all of the gp120 after a 2.5-hr incubation with 10 µM BNM-III-170 (Fig. 4A and B). By contrast, the 130-C and the BNM-III-170-resistant Envs (E64G + I424T, E64G + S375N and E64G + S375N +I424T) were relatively unaffected by incubation with up to 100 µM BNM-III-170. The Env mutants (S375N, I424T, E64G and S375N + I424T) with intermediate levels of BNM-III-170 resistance exhibited gp120 shedding at high doses of BNM-III-170. Thus, there is a strong correlation between the sensitivity of the functional Env variant to BNM-III-170 inhibition of infection and susceptibility to BNM-III-170-induced gp120 shedding from the Env trimer (Spearman rank-order correlation coefficient, r_S_ = −0.97, p<0.05) (Fig. 4C).

**Figure 4.**
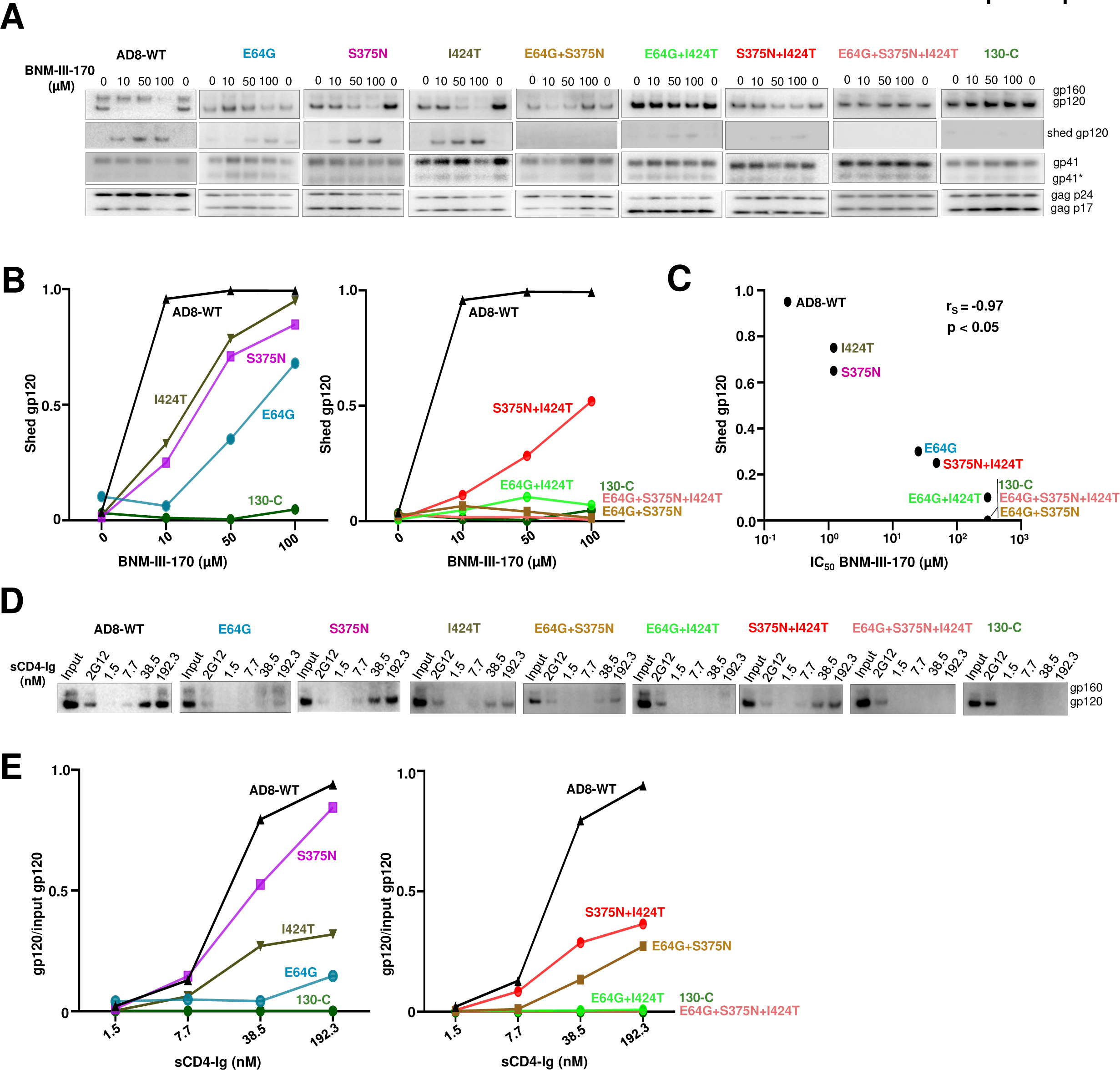
Mechanisms of HIV-1_AD8_ escape from BNM-III-170 and sCD4-Ig. (A) BNM- III-170-induced shedding of gp120 from virions with the indicated Envs was measured. Virions produced transiently from HEK 293T cells transfected with pNL4-3-AD8 proviral constructs were harvested, clarified by low-speed centrifugation, filtered (0.45-µm) and pelleted at 14,000 x g for 1.5 h at 4°C. The virus pellet was resuspended in 1X PBS and incubated with the indicated concentrations of BNM-III-170 for 2.5 h at 37°C. The viruses were then pelleted; the virus pellet was lysed and the supernatant containing shed gp120 was incubated with Galanthus nivalis lectin (GNL) beads. The viral lysates and the protein captured on the GNL beads were Western blotted. (B) gp120 shedding in the presence of BNM-III-170 was measured by quantifying the gp120 bands in the Western blots shown in A. The shed gp120 was calculated according to the formula: gp120 supernatant/ (gp120 supernatant + gp120 virions). (C) The correlation between BNM-III-170 inhibition of pseudovirus infection and BNM-III-170-induced gp120 shedding is shown. The amount of gp120 shedding at a BNM-III-170 concentration of 50 µM, measured as in B, was used in the correlation analysis. The Spearman rank- order correlation coefficient (r_S_) and p value are shown. (D) Precipitation of virion Env by sCD4-Ig was evaluated as an indication of the affinity of sCD4-Ig for the Env variants. Virion pellets prepared as in A were resuspended in buffer containing the indicated concentrations of sCD4-Ig and incubated for 1.5 h at room temperature. After pelleting, the viruses were washed, lysed and incubated with Protein A-agarose beads. The proteins captured on the beads were Western blotted. The results shown are typical of those obtained in at least two independent experiments. (E) The ratio of thevirion gp120 glycoproteins precipitated by sCD4-Ig to the input gp120 glycoproteins was calculated from the experiment shown in D. The ratio is shown as a function of the sCD4-Ig concentration used for the immunoprecipitation assay.

To evaluate the binding of sCD4-Ig to the Env variants, virions with the different Envs were incubated with sCD4-Ig at room temperature; under these conditions, sCD4- Ig-induced gp120 shedding is relatively inefficient (89). The virions were washed and the virion-associated sCD4-Ig was used to precipitate the bound Envs. The sCD4-Ig precipitated the WT HIV-1_AD8_ Env at a half-maximal concentration of approximately 25 nM (Fig. 4D and E). No sCD4-Ig binding was detected to virions with the 130-C Env.

The sCD4-Ig bound to virions with the other Env variants with intermediate levels of efficiency. The E64G change apparently makes a major contribution to the decrease in the efficiency of sCD4-Ig binding observed for the 130-C Env on virions.

### Activation of HIV-1 infection of CD4-negative, CCR5-expressing cells by BNM-III- 170 and sCD4-Ig

We tested the ability of BNM-III-170 and sCD4-Ig to activate infection of CD4-negative, CCR5-expressing cells by the HIV-1_AD8_ Env variants (31, 41, 42, 90, 91). Infection of Cf2Th-CCR5 cells by viruses with WT HIV-1_AD8_ Env was activated by BNM-III-170 and sCD4-Ig (Fig. 5A and B). Activation by BNM-III-170 was less efficient for viruses with the 130-C and other variant Envs. This was also true for sCD4-Ig activation, with one exception; infection of Cf2Th-CCR5 cells by viruses with the S375N + I424T Env was activated by sCD4-Ig more efficiently than infection by viruses with the WT HIV-1_AD8_ Env. The susceptibility of the Env variants to BNM-III-170 and sCD4-Ig activation correlated with their sensitivity to inhibition by these Env ligands (Spearman rank-order correlation coefficients, r_S_ = 0.94 and 0.81, respectively; p<0.05 for both correlations) (Fig. 5C).

**Figure 5.**
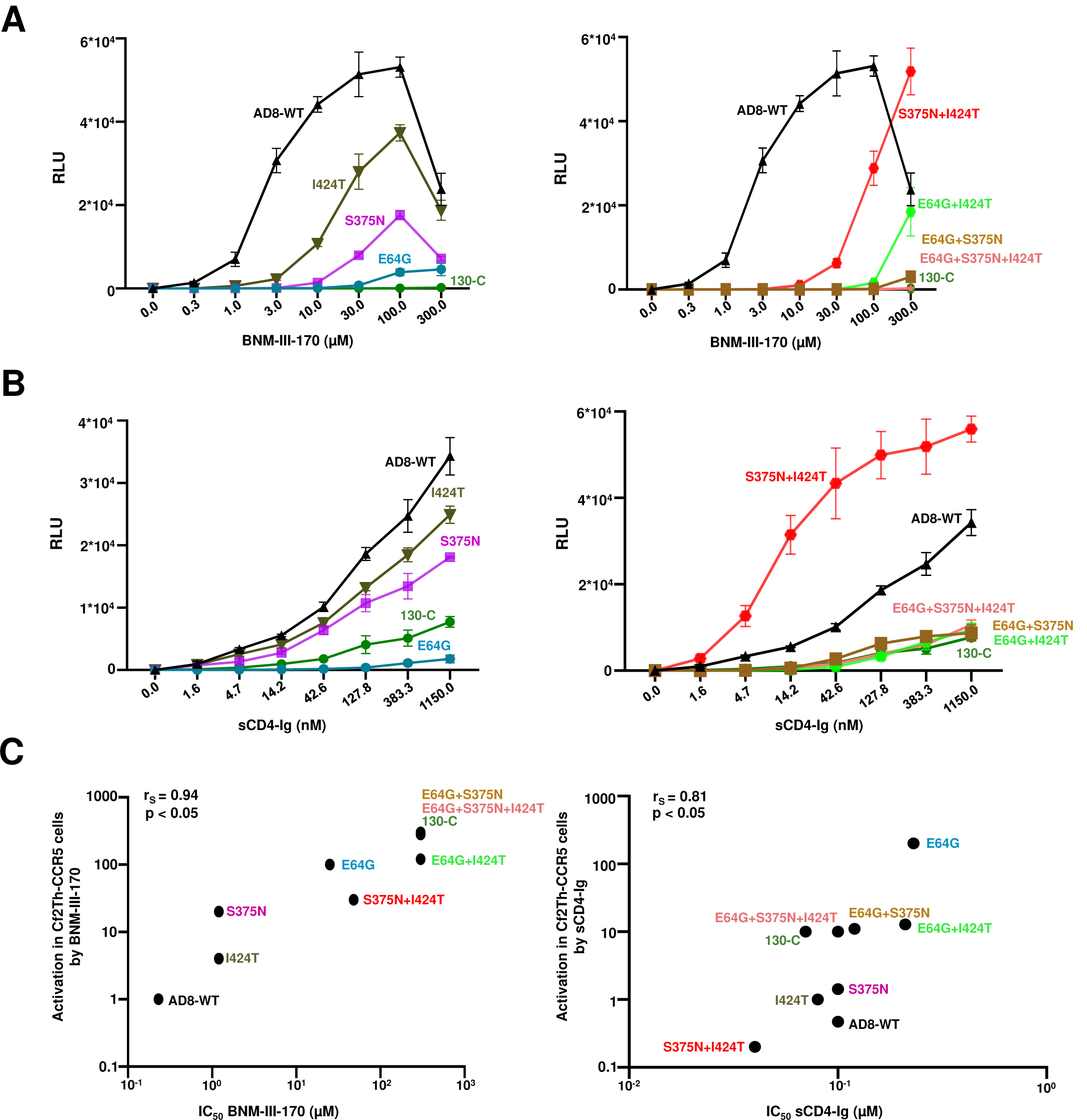
Activation of HIV-1 infection by BNM-III-170 and sCD4-Ig. Activation of HIV-1 infection of CD4-negative, CCR5-expressing cells by BNM-III-170 (A) and sCD4- Ig (B) was evaluated. HEK 293T cells were transfected with plasmids expressing the indicated Envs and HIV-1 packaging proteins and a luciferase-expressing HIV-1 vector. After 48 h, pseudoviruses were harvested and incubated with Cf2Th-CCR5 cells in 96- well plates. The plates were centrifuged at 600 × *g* for 30 min at 21°C. Medium containing serial dilutions of BNM-III-170 or sCD4-Ig was then added. After incubation at 37°C in a CO_2_ incubator for 48 h, the cells were lysed and luciferase activity was measured. RLU, relative light units. (C) The correlation between inhibition of infection of Cf2Th-CD4/CCR5 cells (on the x-axis) and activation of infection of Cf2Th-CCR5 cells (on the y-axis) by BNM-III-170 and sCD4-Ig is shown for the indicated pseudovirus variants. The y-axis values represent the concentrations of BNM-III-170 (in µM) and sCD4-Ig (in nM) required to achieve a level of infection in the Cf2Th-CCR5 cells of 0.5 x10^4^ RLU, determined as in A and B, respectively. The Spearman rank-order correlation coefficients (r_S_) and p values are shown.

The inhibition and activation of the functional Env trimer reflect both the ability of sCD4-Ig to bind Env and to induce concomitant conformational changes. We did not observe a significant correlation between the ability of the virion Envs to bind sCD4-Ig in our assay and the sensitivity of the Env variants to sCD4-Ig inhibition or activation (data not shown). As an example, the S375N + I424T Env bound sCD4-Ig less efficiently than WT HIV-1_AD8_ Env but was inhibited and activated more efficiently than WT HIV- 1_AD8_ Env. Apparently, the assay measuring sCD4-Ig interaction with the virion Env trimer does not capture all of the factors that influence sCD4-Ig inhibition and activation of infection. Likewise, only modest (<4-fold) differences in the affinity of monomeric gp120 from the different Env variants for sCD4 were observed (Table 2), and these did not significantly correlate with sCD4-Ig activation and inhibition of infection by these variants (data not shown).

**Table 2.** Affinity of monomeric gp120 from the different Env variants for sCD4. Association of sCD4 with monomeric gp120 from the different variants was estimated by BLI, with the calculated on-rates (k_a_), off-rates (k_dis_), and dissociation constants (K_D_). E represents 10 raised to the given exponential. The means and standard deviations derived from two independent experiments are shown.

| **gp120** | **K<sub>D</sub> (nM)** | **k<sub>a</sub>(1/Ms)** | **k<sub>dis</sub>(1/s)** |
| --- | --- | --- | --- |
| **AD8-WT** | 13.46 ± 0.05 | 5E+04 ± 5E+03 | 7E-04 ± 7E-05 |
| **E64G** | 11.04 ± 1.93 | 7E+04 ± 2E+04 | 7E-04 ± 7E-05 |
| **S375N** | 17.85 ± 3.63 | 1E+05 ± 2E+03 | 2E-03 ± 3E-04 |
| **I424T** | 4.04 ± 0.02 | 6E+04 ± 3E+03 | 2E-04 ± 1E-05 |
| **E64G+S375N** | 8.83 ± 3.89 | 7E+04 ± 3E+04 | 6E-04 ± 3E-05 |
| **E64G+I424T** | 17.73 ± 6.65 | 1E+05 ± 8E+03 | 2E-03 ± 9E-04 |
| **S375N+I424T** | 25.51 ± 11.10 | 8E+04 ± 2E+04 | 2E-03 ± 3E-04 |
| **E64G+S375N+I424T** | 48.54 ± 1.56 | 6E+04 ± 5E+03 | 3E-03 ± 3E-04 |

### Effects of Env changes on virus sensitivity to conformation-dependent ligands

The sensitivity of HIV-1 infection to inhibition by specific Env ligands or by exposure to cold can provide an indication of global changes in Env conformation (26, 78, 79, 92, 93). Therefore, we examined the susceptibility of the virus variants to inhibition by conformation-sensitive broadly neutralizing antibodies and poorly neutralizing antibodies, the T20 peptide corresponding to the gp41 heptad repeat (HR2) region (94, 95) and cold incubation. The sensitivities of the viruses pseudotyped by the WT HIV- 1_AD8_ and variant Envs to these treatments were similar, with two exceptions (Table 3). Viruses pseudotyped by Envs with the E64G change were approximately 15-fold more sensitive to inhibition by T20 than the other viruses. Viruses with the 130-C Env were slightly less sensitive to cold inhibition than the WT HIV-1_AD8_ or other viruses. These results indicate that BNM-III-170 escape is not accompanied by global changes in Env conformation.

**Table 3.**
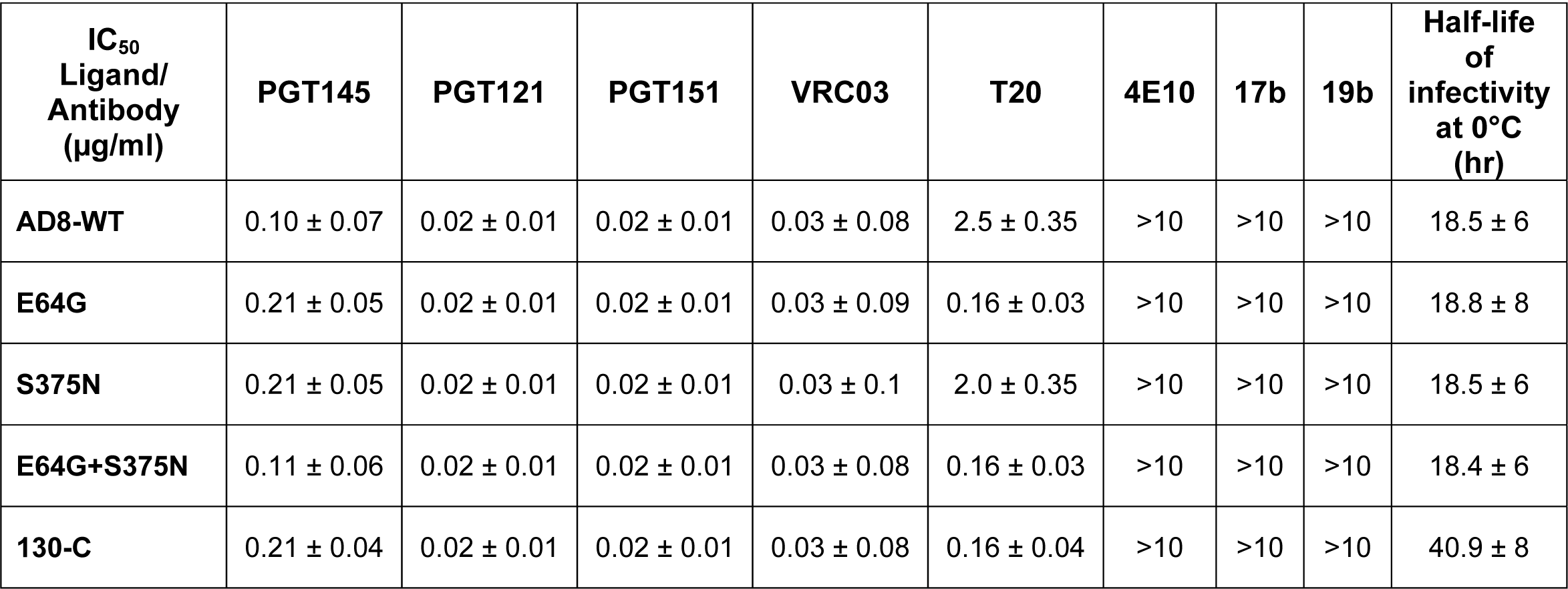
Sensitivity of HIV-1_AD8_ Env variants to antibody neutralization and cold inactivation. The sensitivity of viruses pseudotyped with the indicated HIV-1_AD8_ Env variants to poorly and broadly neutralizing antibodies was evaluated. HEK 293T cells were transfected with plasmids encoding the indicated Envs and HIV-1 packaging proteins as well as a luciferase-expressing HIV-1 vector. Forty-eight h later, cell supernatants were clarified and incubated with different antibodies for 1 hr at 37°C before the mixture was added to Cf2Th-CD4/CCR5 target cells. The concentration of antibody required to inhibit 50% of virus infection (IC_50_) was calculated using the GraphPad Prism program. In the cold sensitivity assay, viruses were incubated on ice for the indicated times, after which the virus infectivity was measured. The results shown are representative of those obtained in at least two independent experiments. The means and standard deviations derived from two independent experiments are shown.

### Susceptibility of Env variants to BNM-III-170 induction of Env epitopes for ADCC

In HIV-1-infected cells, the formation of Env-CD4 complexes potentially exposes CD4- induced (CD4i) Env epitopes that can be recognized by ADCC-mediating antibodies (46–48). These Env-CD4 complexes are down-regulated from the infected cell surface by the HIV-1 Vpu and Nef proteins (55–61). CD4mcs like BNM-III-170 induce conformational changes in HIV-1 Env that expose CD4-induced epitopes on cell-surface Env, which can then be targeted by ADCC responses (50, 53, 54, 62–64). To evaluate the effect of the Env changes on this process, cells infected with molecularly cloned viruses containing the Env variants were incubated with BNM-III-170 or the DMSO control and tested for the ability to be recognized by antibodies. Figure 6 shows that, although the WT HIV-1_AD8_ Env responds to BNM-III-170 by exposing epitopes recognized by the 17b CD4i antibody and antibodies from the plasma of an HIV-1- infected individual, the variant Envs tested did not. The cells with the WT HIV-1_AD8_ Env and variant Envs down-regulated CD4 and BST-2 comparably; this was expected, as these functions are mediated by the viral Vpu and Nef proteins produced by the integrated proviruses (55–61). These results are consistent with the relatively low ability of the 130-C, E64G + S375N and E64G + S375N + I424T Envs to bind and respond to BNM-III-170.

**Figure 6.**
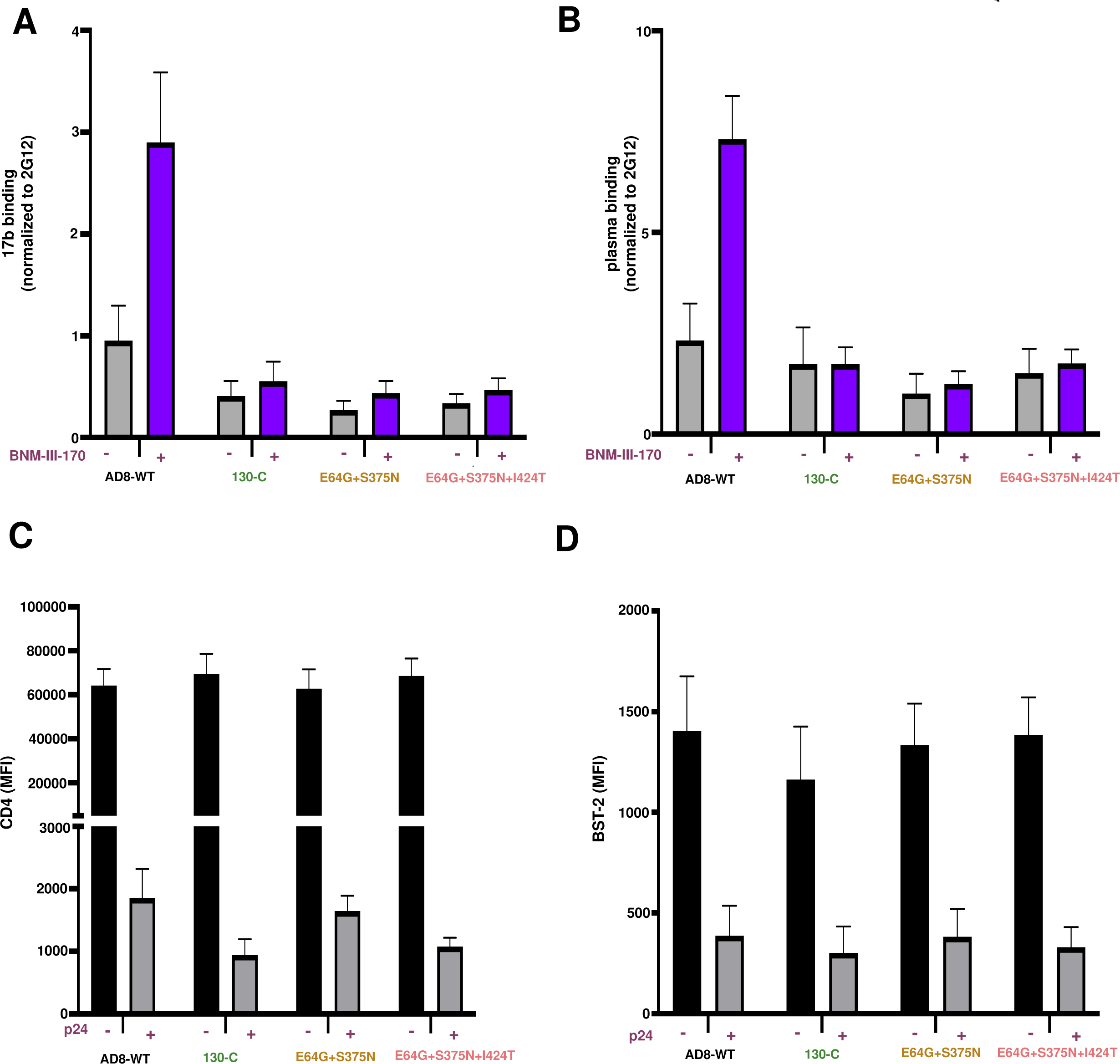
Effect of HIV-1_AD8_ Env changes on BNM-III-170-induced exposure of ADCC epitopes. Activated primary CD4+ T cells were infected with the molecularly cloned viruses expressing the indicated Env variants. Two days after infection, cells were incubated with BNM-III-170 (50 µM) or DMSO and stained with anti-CD4, anti-BST-2 or 17b, 2G12 and A32 antibodies or HIV+ plasma followed by detection of intracellular p24. (A,B) The binding of 17b (A) or HIV+ plasma (B) relative to the binding of 2G12 is shown. Similar results were obtained with the A32 antibody (not shown). (C,D) The mean fluorescence intensity (MFI) for the staining of cells with antibodies against CD4 (C) and BST-2 (D) is shown. Means and standard deviations from two independent experiments are shown.

## DISCUSSION

CD4mcs compete with CD4 for binding to HIV-1 gp120 (40, 42, 91). In the process of binding, CD4mcs induce conformational changes in Env similar to those induced by CD4 (31, 41, 42). These conformational changes ultimately contribute to the binding affinity of the CD4mcs. Thus, resistance to CD4mcs can involve changes in HIV-1 Env near the binding site or changes outside the binding site that decrease the ability of Env to undergo the appropriate conformational changes (35, 67–69, 76, 79).

Selection of HIV-1 variants resistant to CD4mc can provide insights into preferred viral pathways to achieve resistance while minimizing fitness costs. BNM-III-170 is currently the CD4mc with the highest potency and greatest breadth against HIV-1 (37). The viruses selected for resistance to BNM-III-170 exhibited a modest decrease in the ability of their Envs to support virus infection. Most of this fitness cost resulted from the substitution of a glycine residue for the highly conserved Glu 64 in Layer 1 of the gp120 inner domain. Modest decreases in the binding and antiviral potency of sCD4-Ig were associated with the E64G change. Conformational transitions in Layer 1 of gp120 are thought to contribute to the binding of CD4 and CD4mcs, and the E64G change likely decreases the efficiency of this process (78, 80, 87). A large gp120 ligand like CD4 is more able to overcome the resistance to conformational change imposed by E64G than a small molecule like BNM-III-170. Indeed, the E64G change resulted in only 2-3-fold resistance to sCD4-Ig but approximately 100-fold resistance to BNM-III-170. We note that a gp120 region including Glu 64 has been proposed as a second contact site for CD4 on the adjacent Env protomer (96, 97). However, such quaternary contacts, which are not relevant to the binding of a small molecule like a CD4mc, don’t explain the significantly greater impact of the E64G change on susceptibility to BNM-III-170 compared with sCD4-Ig. We also note the proximity of the gp120 inner domain Layer 1 to the gp41 heptad repeat (HR1) region in structural models of Env trimers (34, 77, 82, 98–100). The CD4-induced formation and exposure of the HR1 coiled coil on the gp41 prehairpin intermediate creates a target for antiviral peptides like T20 that mimic the HR2 region of gp41 (94, 95, 101, 102). Slowing Layer 1 conformational transitions may increase the window of exposure of the gp41 HR1 region, accounting for the observed increase in sensitivity of the variants with Gly 64 to T20 inhibition. We did not detect changes in the sensitivity of the BNM-III-170-resistant viruses to neutralization by the antibodies tested, suggesting that resistance is not accompanied by global changes in Env architecture.

The CD4-binding site on gp120 is one of the few conserved protein surfaces exposed on the pretriggered (State-1) Env trimer (30, 103). The associated Phe-43 cavity, although largely inaccessible to protein ligands, allows the binding of small molecules and is well conserved among HIV-1 Group M viruses. Thus, it is no coincidence that multiple small molecules that inhibit the entry of a variety of HIV-1 strains utilize the Phe-43 cavity for binding Env (33-37, 70-72, 76, 77, 82). However, as we observed in this study, HIV-1 can exploit differences in the requirements for CD4 and CD4mc binding by altering residues like Ser 375 or Ile 424 that line the walls of the Phe-43 cavity. The resulting changes in the shape of the Phe-43 cavity exert only modest effects on CD4 binding affinity and are well tolerated with respect to HIV-1 replication. However, the S375N and the I424T changes individually result in approximately 5-fold effects on virus sensitivity to BNM-III-170; when combined, these changes lower the inhibitory potency of BNM-III-170 by approximately 200-fold. Thus, the different dependence of CD4mcs and CD4 binding on the specific shape of the Phe- 43 cavity creates opportunities for HIV-1 to escape the CD4mcs. We note that changes in Ser 375 and other residues lining the Phe-43 cavity have been shown to confer resistance to less potent CD4mcs (35, 44). The gp120 changes that resulted in HIV-1 resistance to the direct antiviral effect of BNM-III-170 also negated the ability of the CD4mc to enhance the binding of antibodies that potentially mediate ADCC lysis of infected cells.

BMS-806 analogues can bind a pretriggered (State-1) conformation of HIV-1 Env and block some of the conformational changes resulting from CD4 binding (19, 32, 83- 86). In the course of binding Env, BMS-806 analogues do not need to induce major conformational changes in Env; therefore, they bind with higher affinity and achieve greater potency than the CD4mcs (12, 15, 31, 34, 82). Nonetheless, as for the CD4mcs, changes in the residues lining the Phe-43 cavity can result in significant HIV-1 resistance to BMS-806 (33, 104, 105). The S375N change had the biggest impact on HIV-1 sensitivity to BMS-806, and the combined S375N + I424T changes resulted in profound levels of resistance. Of interest, changes in Ser 375, including substitution of an asparagine residue, have been observed in resistant HIV-1 variants derived from individuals treated with BMS-806 analogues (104, 105). Like the I424T change, these alterations of the Phe-43 cavity likely affect the accommodation of the BMS-806 benzoyl group (34, 82). Changes in Met 426 and Met 434, which flank the methoxy pyrrolopyridine group residing in the gp120 water-filled channel, have also been associated with resistance in HIV-1-infected individuals treated with these conformational blockers (104, 105). Consistent with the expectation that CD4-induced conformational changes are not required for BMS-806 binding (12, 14, 34, 82), the E64G change had no impact on BMS-806 potency. Despite the opposing mechanisms of Env inhibition by conformational blockers like BMS-806 and CD4mcs, shared elements of their binding sites in the Phe-43 cavity of gp120 create opportunities for HIV-1 to develop resistance simultaneously to both classes of entry inhibitors.

Clinical applications of HIV-1 entry inhibitors targeting the gp120 Phe-43 cavity may be affected by the potential of the virus to develop resistance. Variation of Ser 375 in natural HIV-1 strains can influence the efficacy of CD4mcs, either positively or negatively (73, 76, 106). Viruses with Thr 375 are more susceptible to inhibition by CD4mc than matched viruses with Ser 375 (73, 76). On the other hand, the His 375 residue in Clade AE HIV-1 fills the Phe-43 cavity and renders these viruses resistant to CD4mc (73, 76). Consideration of HIV-1 variation that influences CD4mc efficacy will be important for their successful application in therapy or prophylaxis.

## MATERIALS AND METHODS

### Ethics Statement

Written informed consent was obtained from all study participants and research adhered to the ethical guidelines of CRCHUM and was reviewed and approved by the CRCHUM Institutional Review Board (Ethics Committee, approval number CE 16.164 - CA).

Research adhered to the standards indicated by the Declaration of Helsinki. All participants were adult and provided informed written consent prior to enrollment in accordance with Institutional Review Board approval.

### Cell lines and primary cells

293T cells were grown in Dulbecco’s modified Eagle’s medium (DMEM) (Life Technologies, Wisent Inc.) supplemented with 10% fetal bovine serum (FBS) (Life Technologies, VWR) and 100 μg/ml of penicillin-streptomycin (Life Technologies, Wisent Inc.). Cf2Th cells stably expressing the human CD4 and CCR5 coreceptors for HIV-1 were grown in the same medium supplemented with 0.4 mg/ml of G418 and 0.2 mg/ml of hygromycin. C8166-R5 cells were generously provided by Dr. Seth Pincus, Montana State University (107). The C8166-R5 cells were grown in RPMI 1640 medium (Life Technologies) with 10% FBS. Puromycin (Life Technologies) (1 μg/mL) was used for selection and was added every fifth passage.

Human peripheral blood mononuclear cells (PBMCs) from HIV-negative individuals were obtained by leukapheresis and Ficoll-Paque density gradient isolation and were cryopreserved in liquid nitrogen until use. CD4+ T lymphocytes were purified from resting PBMCs by negative selection using immunomagnetic beads according to the manufacturer’s instructions (StemCell Technologies). The cells were activated with phytohemagglutinin-L (10 µg/mL) for 48 h and then maintained in RPMI 1640 complete medium supplemented with recombinant interleukin-2 (rIL-2) (100 U/mL).

### CD4-mimetic compound

The small-molecule CD4-mimetic compound (CD4mc) BNM-III-170 was synthesized as described previously (108). The compounds were dissolved in dimethyl sulfoxide (DMSO) at a stock concentration of 10 mM and diluted to 50 µM in phosphate-buffered saline (PBS) for cell-surface staining or in RPMI-1640 complete medium for ADCC assays.

### Antibodies and plasma

Antibodies and sCD4-Ig used in neutralization assays are described in reference 109. BMS-806 was purchased from Selleckchem. The anti-coreceptor binding site 17b monoclonal antibody (mAb) (NIH HIV Reagent Program) and plasma from HIV-1-infected individuals were used to assess cell-surface Env conformation. Plasma from HIV-infected individuals was collected, heat-inactivated and conserved at −80 °C until use. The conformation-independent anti-gp120 outer-domain 2G12 mAb (NIH HIV Reagent Program) was used to normalize Env expression. Mouse anti-human CD4 (clone OKT4; Thermo Fisher Scientific) and mouse anti-human BST-2 (clone RS38E, PE-Cy7-conjugated; Biolegend, USA) were also used as primary antibodies for cell-surface staining. Goat anti-mouse IgG (H+L) and goat anti-human IgG (H+L) (Thermo Fisher Scientific) antibodies pre-coupled to Alexa Fluor 647 were used as secondary antibodies in flow cytometry experiments.

### Selection of BNM-III-170-resistant HIV-1_AD8_

293T cells were transfected with the pNL4-3-AD8 provirus construct using effectene transfection reagent (Qiagen). Forty-eight h afterwards, the virus-containing cell medium was collected, clarified by low-speed centrifugation (600 × *g* for 10 min), and filtered through a 0.45 μm membrane. C8166-R5 cells were then incubated with the virus preparation. Twenty-four h later, cells were washed and fresh medium was added. In the first two passages of the cells, the medium was supplemented with 5 μM BNM-III-170. The concentration of BNM-III- 170 in the culture medium was gradually increased with every passage. Virus replication was monitored by a quantitative real-time PCR assay measuring the viral reverse transcriptase activity (110). Briefly, a small amount of cell supernatant was collected and lysed (Lysis buffer: 50mM Tris HCl pH 7.4, 25mM KCl, 0.15% Triton X- 100, 20% glycerol and RNAase inhibitor) as a source of reverse transcriptase. MS2 RNA (Roche Diagnostics) was used as the template. RT PCR master mix and RNAse inhibitor were obtained from Life technologies. The oligos (5’ tcctgctcaacttcctgtcga, 5’ cacaggtcaaacctcctaggaatg and 6[FAM]cgagacgctaccatggctatcgctgtag[TAMsp]) for the reverse transcriptase assay were synthesized by Integrated DNA technologies.

### RNA isolation and cloning

Viral RNA was isolated from the supernatant of BNM-III- 170-treated C8166-R5 cells using the QIAamp viral RNA minikit (Qiagen). cDNA was synthesized using Superscript III reverse transcriptase (Thermo Fisher Scientific) with gene-specific primer (5’ tcgtctcattctttcccttacagcaggccat). Using cDNA as the template, the HIV-1 *env* sequence was amplified using primers (forward primer: 5’ atctagaattcgatatgacaaaagccttaggcatctccta, reverse primer: 5’ tcaggcggccgcttatagcaaaatcctttccaagccct) flanking the Env region and cloned into pCDNA3.1 vector. The amplified *envs* were sequenced to identify changes potentially responsible for BNM-III-170 resistance.

The *env* sequence changes potentially associated with viral resistance to BNM-III-170 were introduced individually or in combination into the pSVIIIenv AD8 plasmid (to produce pseudoviruses in conjunction with a packaging plasmid and HIV-1 vector) or the pNL4-3-AD8 provirus plasmid (to produce infectious virions) using a Q5 site-directed mutagenesis kit (New England BioLabs).

The presence of the desired mutations was confirmed by DNA sequencing.

### Expression and processing of the HIV-1_AD8_ Env variants associated with BNM-III- 170 resistance

HEK 293T cells were transfected transiently with pNL4-3-AD8 proviral plasmids with *env* variants. Forty-eight h later, the cell supernatant was cleared (600 × *g* for 10 min) followed by filtration through a 0.45 μm membrane. The viruses were pelleted by centrifugation at 14,000 x g for 1.5 h at 4°C and lysed. The cell and virus lysates were Western blotted and probed with a goat polyclonal anti-gp120 antibody (Invitrogen), the 4E10 anti-gp41 antibody (NIH HIV Reagent Program) or a rabbit anti-Gag antibody (Abcam). The cell lysates were also Western blotted and probed with a mouse anti-β-actin antibody (Invitrogen).

His_6_-tagged HIV-1_AD8_ gp120 glycoproteins were produced by introducing the His_6_ tag at the gp120-gp41 junction. The gp120 glycoproteins with the BNM-III-170- resistance-associated changes and the corresponding WT HIV-1_AD8_ gp120 glycoprotein were produced transiently in 293T cells and purified by Ni-NTA affinity chromatography.

### Production of recombinant pseudoviruses expressing luciferase

As described previously (26), 293T cells were transfected with pSVIIIenv-AD8 plasmids expressing Env variants, the pCMVΔP1Δenv HIV-1 Gag-Pol packaging construct and the firefly luciferase-expressing HIV-1 vector at a 1:1:3 µg DNA ratio using effectene transfection reagent (Qiagen). Recombinant, luciferase-expressing viruses capable of a single round of replication were released into the cell medium and were harvested 48 h later.

The virus-containing supernatants were clarified by low-speed centrifugation (600 × *g* for 10 min) and used for single-round infections.

### Virus infectivity, neutralization, and cold sensitivity

Single-round virus infection assays were used to measure the ability of the Env variants to support virus entry, as described previously (26). To measure the infectivity of the Env pseudotypes, recombinant viruses were added to Cf2Th target cells expressing CD4 and CCR5.

Forty-eight h later, the target cells were lysed and the luciferase activity was measured.

For neutralization assays, the compounds/antibodies to be tested were incubated with pseudoviruses for 1 h at 37°C. The mixture was then added to Cf2Th target cells expressing CD4 and CCR5. Forty-eight h later, the target cells were lysed and the luciferase activity was measured.

To evaluate the cold sensitivity of the Env variants, pseudotyped recombinant viruses were incubated on ice for various lengths of time prior to measuring their infectivity, as described previously (26, 27, 79, 93).

### gp120 shedding

To measure BNM-III-170-induced gp120 shedding, 293T cells were transfected transiently with pNL4-3-AD8 provirus constructs. The transfected cell supernatants were harvested 48 h later, clarified by low-speed centrifugation (600 × *g* for 10 min) and filtered through a 0.45 μm membrane. The viruses were pelleted by centrifugation at 14,000 x g for 1.5 h at 4°C and resuspended in 100 µl of 1X PBS. The viruses were incubated with different concentrations of BNM-III-170 for 2.5 h at 37°C, followed by centrifugation at 14,000 x g for 1.5 h at 4°C. The virus pellets were lysed 1X LDS buffer and analyzed as described below. The supernatants containing shed gp120 were bound to Galanthus nivalis lectin (GNL) beads (Thermo Fisher Scientific). The glycoproteins captured on the beads and the lysates of virus pellets were Western blotted and probed with a goat anti-gp120 antibody, the 4E10 anti-gp41 antibody or a rabbit anti-Gag antibody. The Western blots were quantified using Image J software (111).

### Immunoprecipitation of virion Envs with sCD4-Ig

Virions produced following transfection with pNL4-3-AD8 proviral plasmids expressing Env variants were pelleted as described above. The pellets were incubated with sCD4-Ig at the indicated concentrations for 1.5 h at 37°C. After washing with 1X PBS, viruses were lysed using buffer (5M NaCl, 1.5M Tris-HCl pH 7.4,1:20 dilution of 10% NP40 and 1X protease inhibitor (Sigma Aldrich)) and incubated with Protein A-agarose beads (Thermo Fisher Scientific). The proteins captured on the beads were Western blotted and probed with a goat anti-gp120 antibody. The Western blots were quantified using Image J software (111).

### Activation of virus infection by BNM-III-170 and sCD4-Ig

Pseudoviruses were incubated with CD4-negative, CCR5-expressing Cf2Th-CCR5 cells in 96-well plates. The plates were centrifuged at 600 × *g* for 30 min at 21°C. Medium containing serial dilutions of sCD4-Ig or BNM-III-170 was then added. Forty-eight h later, cells were lysed, and luciferase activity was measured.

### Biolayer Interferometry

Binding kinetics were performed on an Octet RED96e system (FortéBio) at 25°C with shaking at 1,000 rpm. Amine-Reactive Second-Generation (AR2G) biosensors were hydrated in water, then activated for 300 s with an S-NHS/EDC solution (Fortébio) prior to amine coupling. WT and mutant HIV-1_AD8_ gp120 glycoproteins were loaded into the activated AR2G biosensor at 12.5 µg/mL in 10 mM acetate solution pH 5 (Fortébio) for 600 s and then quenched into 1 M ethanolamine solution pH 8.5 (Fortébio) for 300 s. Baseline equilibration was collected for 120 s in 10X kinetics buffer. Association of sCD4 (in 10X kinetics buffer) to the different gp120 glycoproteins was carried out for 180 s at various concentrations in a two-fold dilution series from 250 nM to 15.625 nM prior to dissociation for 300 s. The data were baseline subtracted prior to fitting, which was performed using a 1:1 binding model and the FortéBio data analysis software. On-rates (k_a_), off-rates (k_dis_), and dissociation constants (K_D_) were computed using a global fit applied to all data.

### Establishment of primary CD4+ T cells infected with Env variants

The vesicular stomatitis virus G (VSV-G)-encoding plasmid was previously described (112). Vesicular stomatitis virus G (VSV-G)-pseudotyped HIV-1 vectors were produced by co-transfection of 293T cells with HIV-1 proviral constructs encoding the variant Envs and a VSV-G-encoding plasmid using the calcium phosphate method. Two days later, cell supernatants were harvested, clarified by low-speed centrifugation (300 × *g* for 5 min), and concentrated by ultracentrifugation at 4°C (100,605 × *g* for 1 h) through a 20% sucrose cushion. Pellets were resuspended in fresh RPMI medium, and aliquots were stored at −80°C until use. Viruses were then used to infect activated primary CD4+ T cells from healthy HIV-1-negative donors by spin infection at 800 × *g* for 1 h in 96-well plates at 25 °C. Viral preparations were titrated directly on primary CD4+ T cells to achieve similar levels of infection among the different proviruses used.

### Flow cytometry analysis of cell-surface staining

Cell-surface staining was performed 48 h after infection of primary CD4+ T cells. Mock-infected or HIV-1-infected primary CD4+ T cells were incubated for 45 min at 37°C with anti-CD4 (0.5 µg/mL), anti-BST-2 (2 µg/mL), anti-Env mAbs (5 µg/mL) or plasma (1:1000 dilution) in the presence or absence of the CD4mc BNM-III-170 (50 µM) or equivalent volume of DMSO. Cells were then washed twice with PBS and stained with the appropriate Alexa Fluor 647- conjugated secondary antibody (2 µg/mL), when needed, for 20 min at room temperature. After two more PBS washes, cells were fixed in a 2% PBS-formaldehyde solution. Infected cells were then permeabilized using the Cytofix/Cytoperm Fixation/ Permeabilization Kit (BD Biosciences) and stained intracellularly using PE-conjugated mouse anti-p24 mAb (clone KC57; Beckman Coulter, Brea, CA, USA; 1:100 dilution). The percentage of infected cells (p24+) was determined by gating on the living cell population according to staining with a viability dye (Aqua Vivid; Thermo Fisher Scientific). Data were acquired on an LSR II cytometer (BD Biosciences), and data analysis was performed using FlowJo v10.5.3 (Tree Star).

### Statistics

The concentrations of antibodies and other inhibitors that inhibit 50% of infection (IC_50_ values) were determined by fitting the data in five-parameter dose-response curves using GraphPad Prism 8. Spearman rank-order correlation coefficients (r_S_) and p values were calculated using VassarStats (113).

## ACKNOWLEDGMENTS

We thank Ms. Elizabeth Carpelan for manuscript preparation. Antibodies against HIV-1 were kindly supplied by Dennis Burton (Scripps), Peter Kwong and John Mascola (Vaccine Research Center NIH), Barton Haynes (Duke University), Hermann Katinger (Polymun), James Robinson (Tulane University), and Marshall Posner (Mount Sinai Medical Center). We thank the NIH HIV Reagent Program for providing additional reagents. We thank Mario Legault from the FRQS AIDS and Infectious Diseases network for cohort coordination and clinical samples.

This work was supported by grants from the National Institutes of Health (grants AI145547, AI124982, AI129017, AI164562, AI150471, AI148379, AI150322 and AI129769), by an HIV Cure Research Grant from Gilead Sciences, and by a gift from the late William F. McCarty-Cooper. This study was partially supported by a Canadian Institutes of Health Research (CIHR) foundation grant #352417, Team grant #422148 to A.F., and a Canada Foundation for Innovation (CFI) grant #41027 to A.F. A.F. is the recipient of a Canada Research Chair on Retroviral Entry #RCHS0235 950-232424.

